# Metabolic dehalogenation of halomethanes by mercury methylators

**DOI:** 10.64898/2026.04.26.720869

**Authors:** Shanquan Wang, Rui Shen, Yi Su, Qihao Li, Ningning Wu, Qihong Lu, Yuxin Sun, Fei Dang, Yue Lu, Laiguo Chen, Deyi Hou, Bi-Xian Mai, Tong Zhang, Rongliang Qiu

## Abstract

Mercury (Hg) methylators play key roles in the global Hg cycle. Nonetheless, the Hg methylators are widely present in Hg-absent environments and the Hg methylation is a cometabolic process without cell growth, which leave the origin and evolution of the Hg methylation as a mystery. Here, we reported the CCl_4_/trihalomethane-to-dihalomethane dehalogenation by model Hg methylators in a metabolic way. Heterologously-expressed HgcAB catalyzed both the CCl_4_ dechlorination and Hg methylation. Halomethanes were shown to sustain the Hg-methylation community without adding external carbon source, electron donor and acceptor. Metadata analyses suggested the halomethane dehalogenation potential of Hg methylators at the global scale. These results, together with much higher global flux and kinetic *V*_max_ of the metabolic halomethanes dehalogenation relative to the co-metabolic Hg methylation, suggested halomethanes as the potential HgcAB substrates for the evolution of Hg methylation.

## INTRODUCTION

Mercury methylators are anaerobic microorganisms to convert inorganic mercury (Hg) into highly toxic methylmercury (MeHg), which play a key role in the global Hg cycle^[1, 2]^. The widespread neurotoxic MeHg can bioaccumulate and magnify up to ten million times throughout the food chain^[3, 4]^, and pose risks to human health and ecosystem safety^[5, 6, 7]^, which in particular has received global increasing attention. A two-gene cluster, consisting of *hgcA* and *hgcB*, encodes two proteins essential for microbial methylation of Hg^[8]^, which has been widely used as a biomarker to identify Hg methylators and to predict Hg methylation potential in natural environments^[9, 10, 11]^. Based on the presence of *hgcAB* genes, a phylogenetically diverse range of microorganisms (e.g., sulfate-reducing bacteria and methanogens) have been identified as Hg methylators from varied environments, particularly the sites without Hg contamination^[9, 12, 13, 14]^. Moreover, Hg cannot induce HgcAB expression in Hg methylators, and the Hg methylation is a co-metabolic process that cannot support the cell growth^[8, 12, 15]^. The non-essential and co-metabolic processes (e.g. Hg methylation) generally do not drive the evolution and maintenance of microbial activities, and microorganisms may even lose their key metabolic traits through changing cultivation conditions^[16]^. For example, sulfate-reducing bacteria as a common *hgcAB*-carrying host can lose sulfate reduction activity upon their syntrophic cultivation for 1,000 generation with methanogens under the cultivation condition without sulfate amendment^[16, 17]^. Therefore, there could be metabolic or essential substrate(s), instead of the Hg as a co-metabolic and non-essential substrate, to drive the origin and evolution of the HgcAB in the *hgcAB*-carrying sulfate-reducing bacteria and other Hg methylators, since the continuing evolution and maintenance of *hgcAB* genes require energy compensation in cells.

Bioinformatic and further structural modeling analyses have indicated that HgcA and HgcB contain an N-terminal cobalamin binding domain and two [4Fe-4S] clusters, respectively^[8, 18]^. The coupling of cobalamin and iron-sulfur clusters, which serve as essential components in Hg methylation, represents a widespread and crucial strategy in nature for driving a range of biotransformation, e.g. carbon-halogen bond cleavage and epoxide reduction^[19]^. Previous studies showed that similar catalytic steps might be shared by the dihalomethane reductive dehalogenase (RDase)-mediated carbon-halogen bond cleavage with the HgcAB-mediated Hg methylation. For instance, in the *mec* (methylene chloride catabolism of dihalomethanes) gene cassette-mediated carbon-halogen cleavage process, methyltransferase MecE catalyzes an initial chloromethyl transfer reaction concomitant with the release of one chlorine substituent from dichloromethane to the corrinoid-binding protein MecB, in which the Fe-S clusters-containing protein MecH serves as a reductive activator for MecB^[20, 21]^. Given the functional similarity of cobalamin and iron-sulfur clusters in both the Hg methylation and carbon-halogen bond cleavage, as well as the co-existence of Hg(II) and halomethanes on the early and modern Earth^[22, 23, 24]^, it is rational to hypothesize that Hg methylators may employ HgcAB to mediate both the halomethane dehalogenation and Hg methylation, and halomethanes can be a potential candidate to drive the origin and evolution of Hg methylation. To test this hypothesis, we selected Hg methylation model strains (*Pseudodesulfovirio mercurii* ND132 and *Geobacter sulfurreducens* PCA) to investigate their halomethane dehalogenation activities that was confirmed with heterologously-expressed HgcAB. Further synthetic microcosm experiments were conducted to test whether halomethanes could sustain the microbial communities for Hg methylation. Finally, the *hgcAB* gene cluster was employed as a biomarker to characterize the taxonomic and global geographic distribution of mercury methylators and their halomethane dehalogenation potential. Our results provide unprecedented insights into the microbially-mediated Hg cycle and the potential driving force for the origin and evolution of microbial mercury methylation.

## RESULTS

### Microbial reductive dechlorination of CCl_4_ supports cell growth of the mercury methylator

To test the hypothesis that mercury methylators might be capable of removing halogens from halogenated methanes, two representative mercury methylators (i.e. *Pseudodesulfovirio mercurii* ND132 and *Geobacter sulfurreducens* PCA^[25, 26]^) were selected to investigate their reductive dehalogenation of carbon tetrachloride (CCl_4_) and bromoform. Neither of these two strains contains known reductive dehalogenase-encoding genes in their genomes^[27, 28]^. Results showed that both ND132 and PCA completely dechlorinated CCl_4_ into chloroform (CF, CHCl_3_) within 20 days, and further partially into dichloromethane (DCM, CH_2_Cl_2_; Fig. 1a; Fig. S1a). In contrast, complete debromination of CF into dibromomethane was observed after 15 days’ incubation in ND132 and PCA cultures (Fig. S1b; Fig. S1c). No dehalogenation activity was detected in abiotic controls (Fig. 1a; Fig. S1) and in ND132 and PCA cultures amended with perchloroethene, 2345-245-chlorobiphenyl (PCB180) or tetrabromobisphenol-A (TBBPA; Fig. S2). In addition, more efficient dehalogenation of halogenated methanes was observed in ND132 cultures than in PCA cultures (Fig. 1a; Fig. S1). Kinetics of the CCl_4_ dechlorination and Hg methylation were further characterized with the ND132/PCA concentrated cells, based on the fact that both reactions followed the Michaelis-Menten kinetics (Fig. 1b; Fig. 1c). Results showed that ND132 exhibited higher values of both kinetic parameters (*K*_*m*_ and *V*_max_) for CCl_4_ dechlorination compared to strain PCA. For instance, in concentrated cell suspensions, the *K*_*m*_ values were 22.09 µM for ND132 and 2.60 µM for PCA, while their *V*_max_ values were 5.59 and 0.89 µM/h, respectively (Fig. 1b). Notably, the apparent kinetic parameters of CCl_4_ dechlorination had much higher values than these of Hg methylation, e.g., the maximum rates (*V*_max_) of CCl_4_ dechlorination were 952.3- and 526.6-times of Hg methylation in ND132 and PCA concentrated cells, respectively (Fig. 1b; Fig. 1c). These results suggested the apparently higher halomethane dechlorination activity compared to the Hg methylation in mercury methylators.

**Figure 1.**
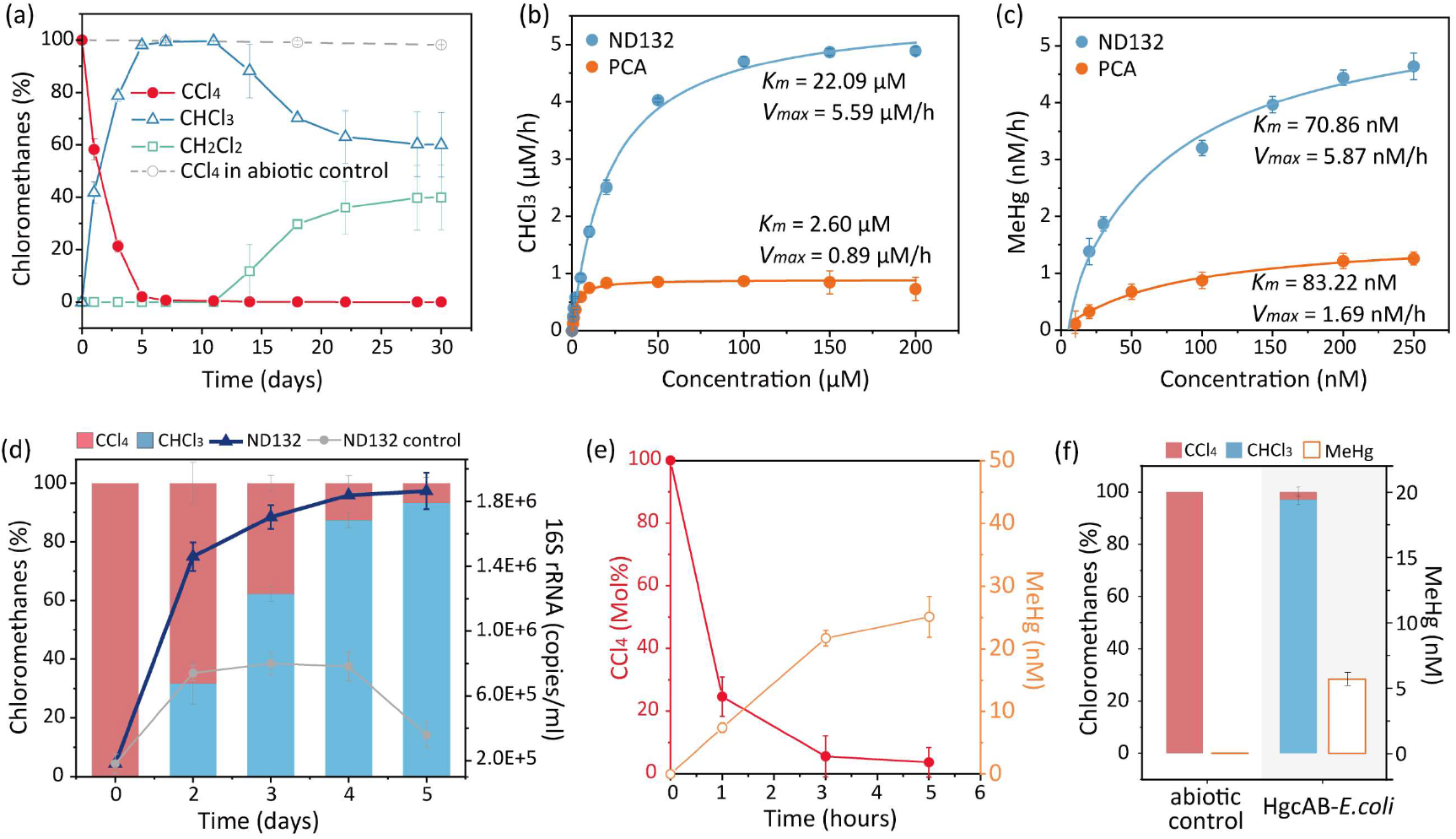
Reductive dechlorination of CCl_4_ by mercury methylators. (a) CCl_4_dechlorination in pre-grown pure culture of *P. mercurii* ND132 and abiotic control without ND132 inoculation. (b) Initial rates of CCl_4_ dechlorination in ND132/PCA concentrated cells with increased CCl_4_ concentrations. (c) Initial rates of Hg methylation in ND132/PCA concentrated cells with increased Hg(II) concentrations. Data points in (b) and (c) were nonlinearly fitted using the Michaelis-Menten equation. (d) Cell growth of ND132 in cultures amended with and without CCl_4_ as a sole electron acceptor. (e) Kinetics of CCl_4_ dechlorination and Hg methylation in ND132 concentrated cells amended with 200 nM CCl_4_ and 200 nM HgCl_2_. (f) Hg methylation and CCl_4_ dechlorination assays with heterologously expressed HgcAB in *E. coli* BL21.

To confirm whether mercury methylators could grow on halogenated methanes, ND132 was selected to monitor its cell growth in the culture amended with CCl_4_ (0.2 µM) as a sole electron acceptor. Results showed the significant microbial proliferation in the CCl_4_-amended ND132 cultures, and the 16S rRNA gene copy number of ND132 increased from 1.81×10^5^ copies per mL on day-0 to 1.86×10^6^ copies per mL on day-5, which was 5.3 times of the concentrations in ND132 control cultures without CCl_4_ amendment (Fig. 1d). The cell growth in the CCl_4_-fed ND132 was 1.4×10^15^ cell per mole of chlorine removed, being around 5 times of the cell growth of PCB-fed *Dehalococcoides mccartyi* CG4^[29]^. These results suggested that ND132 was capable of dechlorinating CCl_4_ in a metabolic way. To compare the substrate preference in the case of CCl_4_-Hg(II) co-existence, ND132 concentrated cells were amended with both CCl_4_ and HgCl_2_ (Fig. 1e). Results demonstrated simultaneous CCl_4_ dechlorination and Hg methylation (Fig. 1e). The heterologously expressed HgcAB in *E. coli* BL21 was shown to reductively dechlorinate CCl_4_ and to methylate Hg (Fig. 1f). In all, these results indicated that mercury methylators could employ HgcAB to metabolically dechlorinate CCl_4_.

### Chloromethane dechlorination-and-degradation derived energy and carbon sources to sustain the microbial consortium for Hg methylation

CCl_4_ dechlorination-derived DCM was reported to be metabolized by a phylogenetically diverse microorganisms (e.g. *Dehalobacterium formicoaceticum* and *Candidatus* Dichloromethanomonas elyunquensis) to form acetate, formate and/or H_2_^[20, 30, 31]^, which can be further employed as both carbon sources and electron donors for the CCl_4_-dechlorination- and Hg-methylation-mediated microorganisms. Therefore, we hypothesize that CCl_4_ can be both energy and carbon sources to sustain a microbial consortium for the Hg methylation without requiring external carbon and energy inputs in natural environments. To test this hypothesis, both the *P. mercurii* ND132 pure culture and a DCM-fermenting microcosm (containing DCM-fermenting *Dehalobacterium*)^[31, 32]^ were used to synthesize a microbial consortium for the Hg methylation and complete dechlorination-and-degradation of CCl_4_, of which the microbial community assembly could be sustained by CCl_4_ amendment (Fig.2).

**Figure 2.**
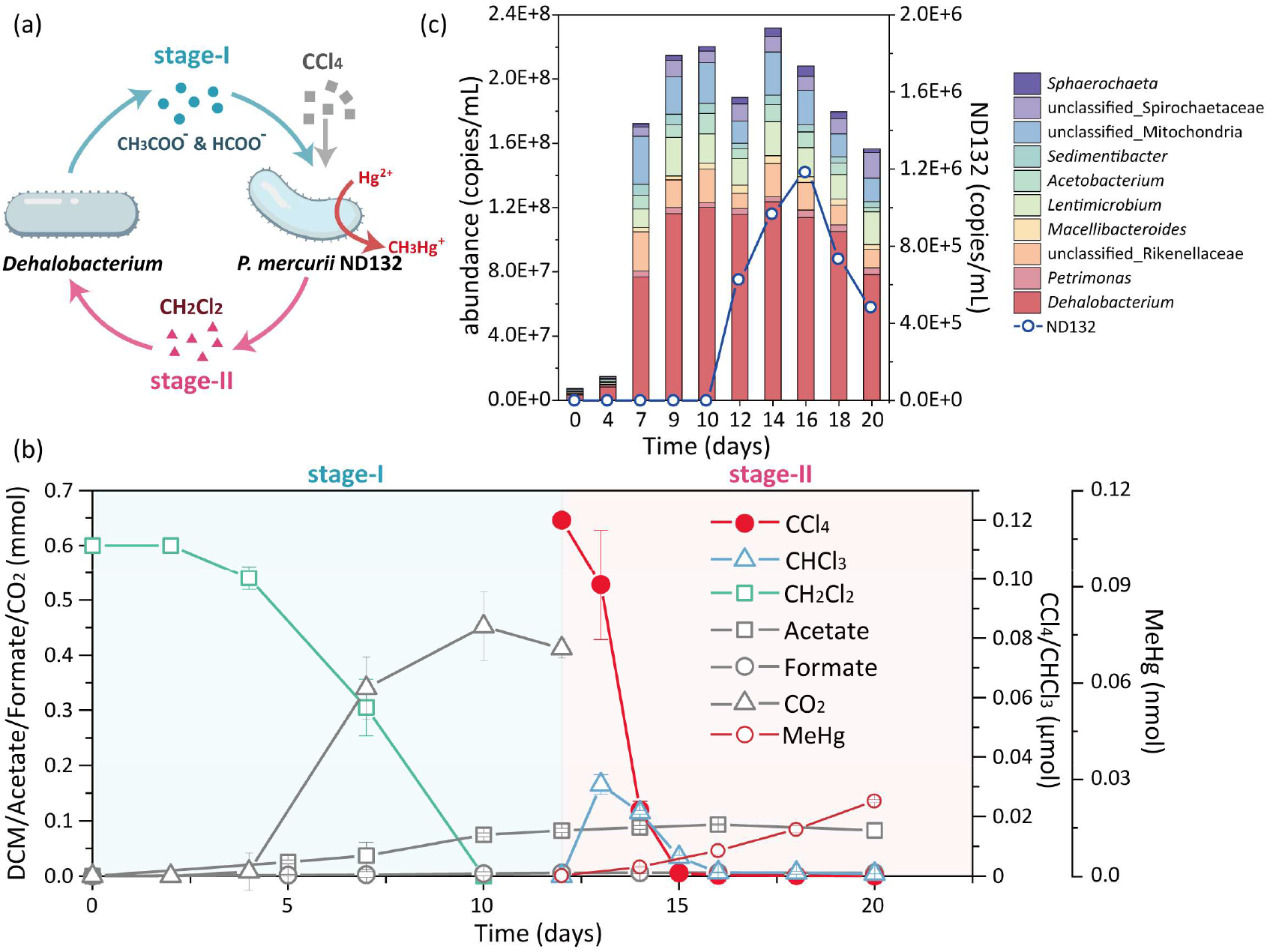
Chloromethanes-driving and -sustaining the microbial community assembling for Hg methylation. (a) Schematic diagram of the two-stage cycling and CCl_4_-sustained microbial community for Hg methylation. (b) Kinetics of chloromethanes dechlorination/degradation, Hg methylation and associated metabolites in the synthetic microbial consortium. (c) Temporal changes in microbial community composition at the genus level (top 10 populations and ND132) based on their absolute abundance in the synthetic microbial consortium. Error bars represent SDs of triplicate cultures.

The incubation experiments proceeded in two-stage cycles (Fig. 2a): stage-I, DCM-degradation to supply acetate and formate for the CCl_4_ dechlorination and Hg methylation; stage-II, ND132-mediated CCl_4_ dechlorination and Hg methylation to provide DCM for subsequent DCM-fermentation and CCl_4_ dechlorination cycles. In the stage-I, 0.60 mmol DCM was completely degraded within 10 days with a maximum amount of accumulated 0.08 mmol acetate and 0.006 mmol formate (Fig. 2b). Accordingly, the total 16S rRNA gene copy number increased from 9.29×10^6^ to 2.45×10^8^ copies per mL (Fig. 2c), in contrast to no microbial growth in control cultures without DCM amendment (Fig. S3). After complete DCM degradation, the 16S rRNA gene copy number of DCM-fermenting *Dehalobacterium* increased from 3.40×10^6^ (on day-0) to 1.20×10^8^ copies per mL (on day-10; Fig. 2c). The methylene chloride catabolism (*mec*) gene cassette was identified as key functional genes for DCM degradation, of which *mecE* as the biomarker gene increased from 8.99×10^5^ to 1.11×10^8^ copies per mL (Fig. S4). In the stage-II with ND132 inoculation and Hg(II)/CCl_4_ amendment, CCl_4_ was completely dechlorinated within three days with a transient accumulation of chloroform (CHCl_3_, a peak amount of 0.03 μmol) (Fig. 2b). Chloroform was further completely dechlorinated and degraded without observing DCM accumulation (Fig. 2b), suggesting the immediate DCM scavenging by *Dehalobacterium*. All chloromethanes in the synthesized microbial consortium were completely removed within 8 days. Notably, the Hg methylation and cell growth of ND132 were observed to co-occur with the chloromethanes dechlorination (Fig. 2b and c). No chloromethanes dechlorination/degradation or Hg methylation was observed in the abiotic control cultures (Fig. S5). Collectively, CCl_4_ and its dechlorination-derived DCM were identified to be capable of driving and sustaining the microbial community assembly for Hg methylation, which indicated the potential critical role of CCl_4_ in the microbially mediated global Hg cycling.

### Taxonomic and geographic distribution of mercury methylators and their correlation with halomethanes in global environments

To evaluate the taxonomic and geographic distribution of mercury methylators, the *hgcAB* genes were screened with global marine and terrestrial metagenomes (totally 2786 metagenomes; Supplementary Data 3; Supplementary Data 4). There was a total of 44,400 *hgcAB*-containing contigs assembled from 954 (out of 1410, 67.66%) marine and 858 (out of 1376, 62.35%) terrestrial samples (Fig. 3), suggesting the widespread presence of mercury methylators. Taxonomic profiling revealed the phylogenetically diverse *hgcAB*-carrying bacteria and archaea across 125 and 10 phyla, respectively (Fig. 3a). Among them, 42.62% *hgcAB*-carrying populations were of Desulfobacterota, Chloroflexota, Acidobacteria, and Bacteroidota (Fig. 3a). Particularly, Dehalococcoidales of the phylum Chloroflexota emerged as a primary *hgcAB*-carrying lineage in marine environments (Fig. S6). Notably, in contrast to the observation that populations of the Dehalococcoidales as typical organohalide-respiring bacteria in terrestrial environments, Dehalococcoidales lose RDase-based organohalide respiration activity in marine environments^[33]^. The widespread presence of *hgcAB* in marine Dehalococcoidales implied that Dehalococcoidales might employ HgcAB in organohalide metabolism in the ocean (Fig. S6a).

**Figure 3.**
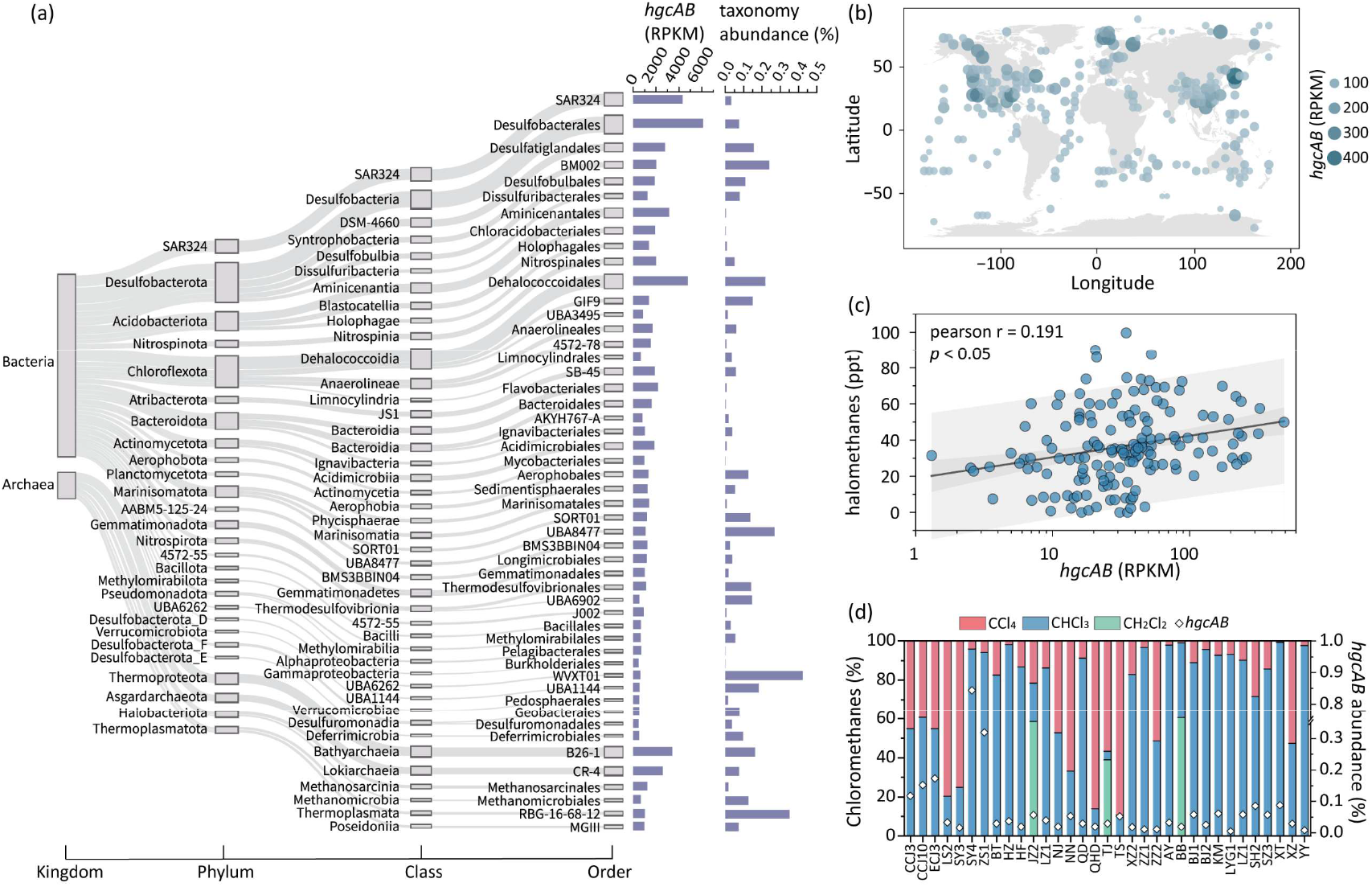
Taxonomic and geographic distribution of mercury methylators and their correlation with halomethane dehalogenation. (a) Major taxonomic distribution of *hgcAB* genes in prokaryotes. (b) Global geographic distribution of *hgcAB* genes in marine and terrestrial environments. Values represent the average abundance of *hgcAB* genes within each 5°×5° (longitude × latitude) grid cell. (c) Correlation analysis between the abundance of *hgcAB* genes and the concentration of halomethanes. (d) Reductive dechlorination of CCl_4_ in representative *hgcAB*-harboring microcosms after 7 days incubation.

For the geographic distribution, *hgcAB* genes were present in diverse geographic locations from Arctic Sea to the Southern Hemisphere (Fig. 3b). There were 68.15% of total *hgcAB* present in marine environments, especially in the marine sediments (55.11% of total *hgcAB*; Fig. S7a). In contrast, terrestrial *hgcAB* genes were evenly distributed, with slightly elevated presence in forest, tundra and wetland samples (Fig. S7b). In addition, the average relative abundance of *hgcAB* genes in the Northern Hemisphere was around two folds of the Southern Hemisphere (78.37 RPKM *vs*. 35.65 RPKM), particularly the coastal regions of the western Pacific Ocean, the eastern North Pacific and the Gulf of Mexico (Fig. 3b). To further correlate the halomethane dehalogenation and mercury methylators, data on global atmospheric concentrations of halomethanes were collected to show their wide distribution with concentrations ranging from 0.76 to 99.65 ppt (an average value of 36.56 ppt; Fig. S8). The CF and DCM as dominant halomethanes and potential dechlorination products had a maximum concentration of 37.64 ppt and 72.54 ppt, respectively (Fig. S9). A significant positive linear correlation (r = 0.191, *p*< 0.05) of *hgcAB* genes with halomethanes was observed to highlight the potential role of mercury methylators in mediation of halomethane dehalogenation in global natural environments. To further confirm the potential role of HgcAB in halomethane dehalogenation, a total of 33 representative marine- and terrestrial samples carrying the *hgcAB* genes were selected to inoculate microcosm cultures to test their activity in reductive dechlorination of CCl_4_. Results showed that all of the *hgcAB* genes-harboring microcosms could dechlorinate 10.77-99.30% CCl_4_ with CF as the dominant dechlorination product after 7 days incubation (Fig. 3d). No obvious CCl_4_ dechlorination was observed in their abiotic controls (Fig. S10). Collectively, these results demonstrate that highly halogenated halomethanes serve as potential substrates to sustain the widespread occurrence of mercury methylators in global natural environments.

## Discussion

Hg methylators employ HgcAB to reductively dehalogenate halomethanes in a metabolic way, in which energy can be harvest through organohalide respiration with halomethanes as electron acceptors. Previous studies have identified the global distribution of *hgcAB* genes in a wide range of environmental habitats where Hg is absent^[9, 12, 13]^, suggesting that the HgcAB may serve physiological functions besides the Hg methylation. Our results provide the empirical example for an additional, or even a cornerstone of, physiological functions of the HgcAB. The HgcAB-catalyzed Hg methylation and halomethane dehalogenation enable the coupling of Hg- and organohalide-cycles in natural environments, which can have profound impacts on the global Hg cycle, and on the evolution of Hg methylation activity based on following facts: (1) over than 65 times of global flux of halomethanes (768 Gg/y CCl_4_, CHCl_3_ and CHBr_3_) relative to inorganic and organic Hg (11.8 Gg/y)^[34, 35]^, and the ratio of estimated global flux of halomethanes to the flux of MeHg is around 1000^[36, 37, 38, 39, 40, 41, 42]^; (2) 1,000 times of kinetic *V*_max_ in the halomethane dehalogenation (0.89-5.59 µM/h) compared to the Hg methylation (1.69-5.87 nM/h), which perfectly matched the global flux ratio of halomethanes to MeHg. These facts well support the hypothesis that Hg methylators in natural environments employ HgcAB to catalyze the halomethane dehalogenation and Hg methylation at the global scale.

The origin and evolution of microbial Hg methylation has been a long-standing mystery. Recently, based on the genome-resolved phylogenetic analyses, Hg methylation was proposed to be evolved for Hg methylators to produce antimicrobial MeHg on a potentially resource-limited early Earth^[43]^. On the early Earth, volcanoes emitted abundant inorganic Hg and reactive halogens (Cl and Br), which were subsequently converted into Hg(II) and organohalides (mainly halomethanes)^[22, 23, 24]^. Both the Hg(II) and halomethanes provided the substrates to drive the origin and evolution of the HgcAB in Hg methylators. Our experiments on the CCl_4_-sustained Hg methylation microcosm, together with the presence of abundant halomethanes on the early Earth and the high kinetic *V*_max_ of the metabolic CCl_4_ dechlorination, support another possibility for the origin and evolution of the HgcAB and associated Hg methylation process (Fig. 4), i.e. on the resource-limited early Earth, the highly volatile, widespread and abundant halomethanes could be the driving force for the origin and evolution of the HgcAB in Hg methylators. Nonetheless, this possibility warrants future extensive molecular investigations to provide empirical examples.

**Figure 4.**
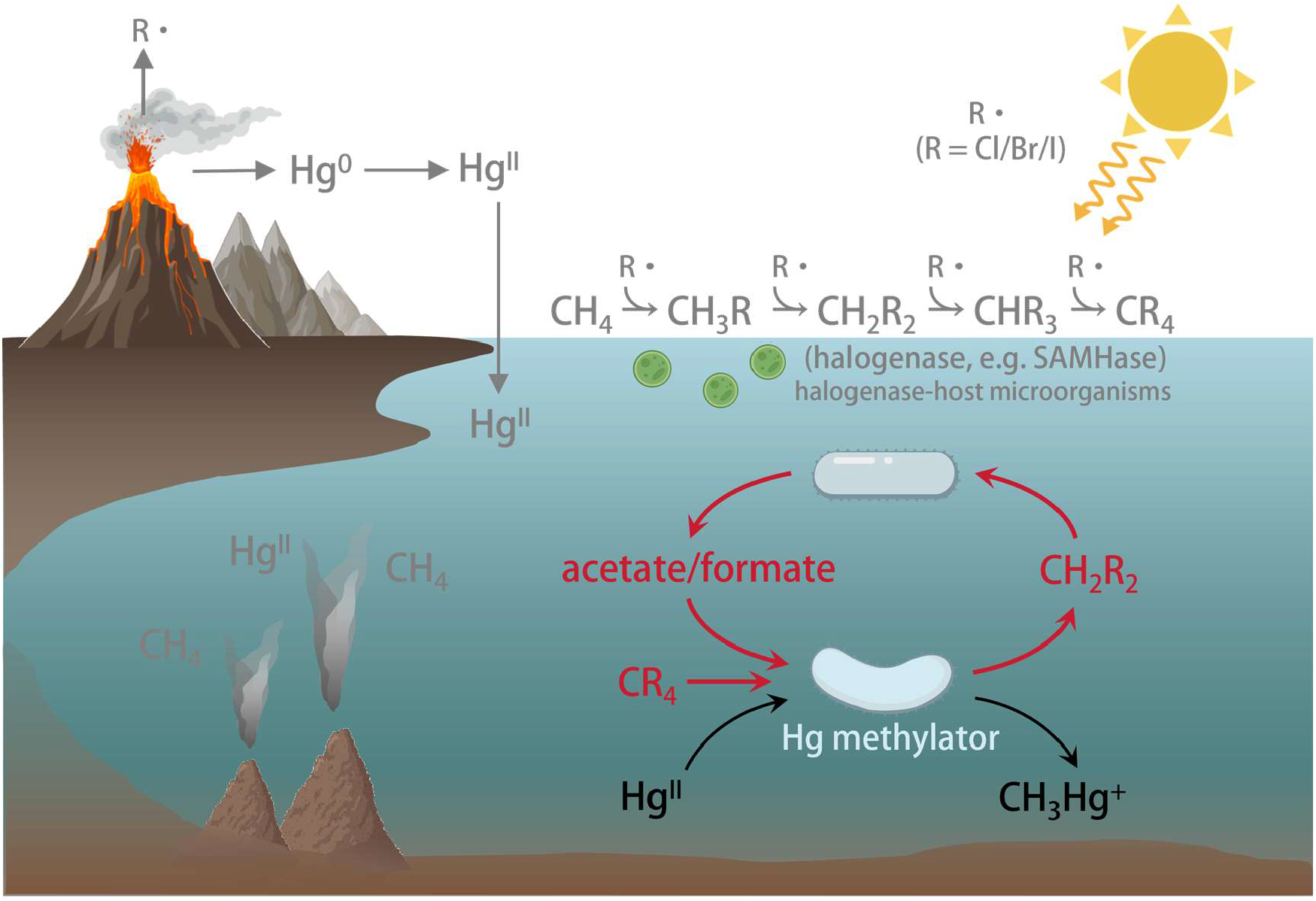
Schematic diagram of the halomethane-supported Hg-methylating microbial community on the early Earth. R·, reactive halogen species; CR_4_, tetrahalomethane; SAMHase, S-adenosyl-L-methionine-dependent halogenase.

For the RDase-catalyzed dehalogenation in organohalide-respiring bacteria (OHRB), a diverse range of OHRB and RDases have been identified from varied environmental sources for the halogen removal from halogenated-ethenes, -ethanes and -aromatics^[33, 44]^. Nonetheless, the enzyme(s) catalyzing the CCl_4_ dehalogenation remain unknown, and previous studies mainly focus on searching the potential CCl_4_-dehalogenation enzymes from traditional RDases. Our study showed the efficient dehalogenation of CCl_4_ and trihalomethanes by Hg methylators without harboring traditional RDase genes, which together with heterologously expressed HgcAB to catalyze CCl_4_ dechlorination and Hg methylation, suggested the key role of the HgcAB in CCl_4_ dehalogenation in natural environments. Similar with MecB/MecH for dichloromethane dechlorination, the HgcA and HgcB for the CCl_4_-to-dichloromethane dechlorination contain cobalamin and iron-sulfur clusters, respectively, which are different from traditional RDases harboring both cobalamin and iron-sulfur clusters in a single protein (RDaseA)^[20, 21, 45, 46]^. In addition, the Dehalococcoidales from terrestrial sources (particularly the organohalide-polluted sites) harbor RDase-encoding genes and are obligate organohalide-respiring bacteria that only employ organohalides as the electron acceptor for the energy metabolism and cell growth^[47, 48]^. Nonetheless, previous studies have identified that marine Dehalococcoidales genomes contain no RDase genes^[33, 49]^, which leave a mystery how the Dehalococcoidales evolve to obtain dehalogenation activity when moving from marine environments to terrestrial niches. The identification of Dehalococcoidales as the major *hgcAB* gene host in the ocean support the hypothesis that marine Dehalococcoidales may employ HgcAB to metabolically dehalogenate halomethanes and later evolve to gain RDases to remove halogens from diverse organohalides under high organohalide stress in terrestrial environments. Though the hypothesis awaits extra experimental evidences, halomethanes as the metabolic substrates of the Hg methylators could spark the global Hg methylation by supporting cell growth of the Hg methylators (especially at co-contaminated sites by Hg and halomethanes), due to the massive generation and widespread of the highly volatile halomethanes at the global scale. These results also hinted that the MeHg-derived ecological risks might be largely underestimated at these co-contaminated sites. Therefore, more intensive and extensive studies are required in future to elucidate the exact impact of the halomethanes on the global Hg methylation.

## Methods and Materials

### Cultivation and medium preparation

The Hg-methylating strains, *Pseudodesulfovirio mercurii* ND132 and *Geobacter sulfurreducens* PCA, were provided by Zhang’s and Lu’s Labs, respectively^[25, 50, 51, 52, 53]^. A representative dehalogenating strain, *Dehalococcoides mccartyi* CG1, was obtained from our previous studies^[29]^. The DCM-fermenting culture was provided by Yang’s Lab^[54]^. These four cultures were cultivated in anaerobic mineral salt medium with tailored carbon sources, electron donors and electron acceptors. Information on the detailed medium compositions was summarized in the Supplementary Data 1. Specifically, *P. mercurii* ND132 was cultured in salt medium amended with sulfate (10 mM) as the electron acceptor and lactate (20 mM) as the carbon source and reducing equivalent^[55]^. *G. sulfurreducens* PCA was cultivated with 20 mM acetate and 20 mM fumarate^[51]^. The *D. mccartyi* CG1 was fed with acetate (10 mM), hydrogen gas (5 × 10^4^ Pa) and PCE (0.5 mM) as a carbon source, electron donor and acceptor, respectively^[29]^. The DCM-degrading culture was cultivated in the salt medium amended with 10 mM DCM without other carbon source and electron donor/acceptor^[56]^. All cultures were incubated at 30°C in the dark without shaking.

### Dehalogenation assays in ND132 and PCA

*P. mercurii* ND132, *G. sulfurreducens* PCA and *D. mccartyi* CG1 were first pre-cultured in above-described salt medium, and then different organohalides were added separately to the late exponential phase cultures, i.e., CCl_4_ (20 μM), bromoform (20 μM), perchloroethene (PCE, 0.5 mM), tetrabromobisphenol A (TBBPA, 5 ppm), and 2345-245-chlorobiphenyl (PCB180, 0.5 ppm). Abiotic controls were prepared without inoculation. Cultures were incubated at 30°C in the dark without shaking.

### Kinetics analysis of CCl_4_ dechlorination and Hg methylation in pure cultures

*P. mercurii* ND132 and *G. sulfurreducens* PCA were pre-cultured in their respective salt medium. At the late exponential phase, cells of *P. mercurii* ND132 and *G. sulfurreducens* PCA were harvested, respectively, and washed three times by repeated centrifugation (1200 ×g, 10 min, 4 °C), followed by resuspension in a deoxygenated phosphate-buffered saline (PBS, pH=7.4) to obtain optical density (OD_600_) values of 2.0. The PBS buffer was first deoxygenated by boiling and purging with ultra-high purity N_2_ gas for > 3 h and then autoclaved. For the substrate concentration experiments, concentrated cells were spiked with Hg(II) (added as HgCl_2_) to final concentrations of 10, 20, 30, 50, 100, 150, 200, 250 nM or with CCl_4_ to final concentrations of 0.5, 1, 2, 5, 10, 20, 50, 100, 150, 200 μM. For substrate preference comparison experiments, concentrated cells were amended with Hg(II) (200 nM) and CCl_4_ (200 nM).

### Cell growth determination

To confirm whether ND132 could grow on CCl_4_, ND132 was sequentially transferred three times in medium amended with lactate (20 mM) as sole electron donor and CCl_4_ (0.2 µM) as sole electron acceptor, followed by an additional transfer with which ND132 cells were collected and quantified. A biotic control without CCl_4_ addition was prepared with the ND132.

### Hg methylation and CCl_4_ dechlorination assays with heterologously expressed HgcAB in *E. coli* BL21

Anaerobic expression of HgcAB in *E. coli* BL21(DE3) was performed as described previously^[57]^. Briefly, the *hgcA* and *hgcB* genes from ND132 were cloned in tandem into pETDuet-1. The resulting plasmid was co-transformed with pRKISC (for [4Fe-4S] cluster assembly) and pBtu (for cobalamin uptake) into *E. coli* BL21(DE3). For expression, cells were grown in M9-ETA medium containing ampicillin (100 mg/L), spectinomycin (50 mg/L) and tetracycline (10 mg/L) under aerobic conditions until OD_600_ = 0.35, then induced with 0.2% (w/v) L-arabinose. At OD_600_ = 0.9, the culture was sparged with N_2_ for 1 h, and 24 mg/L β-D-thiogalactoside (IPTG), 0.36 g/L sodium dithionite, 0.36 g/L cysteine, 50 mg/L ampicillin and 0.2 g/L ammonium iron sulfate were added. Anaerobic expression proceeded for 20 h at 37 °C. After induction of HgcAB, the BL21(DE3) cells were washed and resuspended in deoxygenated PBS solution, using the same parameters as described above for the treatment of ND132 and PCA. After resuspension, 20 nM HgCl_2_ and CCl_4_ were added separately. The cultures were then incubated for 7 days prior to measurement.

### Construction of the halomethanes-sustained microbial consortium

A synthetic consortium was prepared with the ND132 and DCM-degrading culture in mineral salt medium supplemented with 1 mM 2-bromoethanesulfonate (BES) to inhibit methanogenic consumption of acetate and formate^[30]^. The DCM-degrading culture was first inoculated at 1% (v/v) into the medium containing 10 mM DCM. Following the complete depletion of DCM, ND132 that had been washed and resuspended in PBS was inoculated (1‰, v/v), along with the addition of 2000 nM CCl_4_ and 200 nM HgCl_2_. Controls included an abiotic control (sterile medium) and biotic control without DCM or CCl_4_ addition.

### Environmental sample collection and microcosm experiments

Totally 33 representative environmental samples (i.e. 7 marine sediment, 14 urban river sediment and 12 landfill leachates samples) were selected as described previously^[33, 48, 58]^ (Supplementary Data 2). 60 mL of an autoclaved mineral salt medium amended with 10 mM lactate was dispensed into 100 mL serum bottles containing ∼0.1 g of sediment samples or 1 mL of leachate samples. Then microcosms were spiked with 2 μM CCl_4_ as an electron acceptor. For abiotic controls, the sediment- or leachate-amended microcosms were autoclaved prior to CCl_4_ amendment.

### DNA extraction, Sequencing, and analyses

Cells for DNA extraction were harvested from 1 mL cultures by centrifugation (15 min, 10,000 × g, 4 °C). The FastDNA Spin Kit for Soil (MP Biomedicals, Carlsbad, CA, USA) was used to extract genomic DNA according to the manufacturer’s instructions. Cell quantification in microbial communities based on high-throughput sequencing was performed as described previously ^[58]^. To quantify ND132 cell proliferation in the CCl_4_-amended cultures, strain-specific primer set 78F(5′-CTG AAG ATC GGG ATA ACC GC-3′)/234R(5′-TTG CCT TGG TAG GCC ATT AC-3′) was used for qPCR enumeration of ND132. The primer set 828F(5′-ACC ATA TTG TCT TTT TGC CYC AG-3′)/1007R(5′-TAC CGC CCA AAT TTY TCT GC-3′) was used for qPCR enumeration of *mecE* gene^[21]^. The qPCR (CFX96 Touch System; Bio-Rad, CA, USA) was performed with QuantiTect SYBR Green PCR kit. For microbial community analysis, the 16S rRNA gene was amplified with the U515F forward primer (5’-GTG CCA GCM GCC GCG GTA A-3’) and U909R reverse primer (5’-GGC CCC GYC AAT TCM TTT RAG T-3’) as described ^[59]^. To pool multiple samples in one Illumina sequencing run, a specific 8-mer barcode for each sample was added to the 5’ end of both forward and reverse primers. PCR amplification and purification were performed as described previously ^[33]^. Purified PCR products were pooled and sequenced using Illumina MiSeq (PE250) sequencing by GENEWIZ (Suzhou, China). Paired-end reads (2 × 250 bp) were processed to generate amplicon sequence variants (ASVs) using the DADA2 (v1.6)^[60]^ package in R (v4.3.2). ASVs were processed using UPARSE. ASVs taxonomies were assigned using RDP classifier with a confidence cut-off of 80% (SILVA, v132).

### Analytical Methods

CCl_4_ and bromoform, as well as their dehalogenation products were analyzed using Agilent 7890B gas chromatograph (GC) equipped with an electron capture detector (ECD), DB-5 capillary column (30 m × 0.32 mm × 0.25 μm, Agilent J&W Scientific, Folsom, CA, USA). The temperature program was initially held at 45 °C for 1 min, increased at 30 °C/min to 150 °C, and held for 2 min. Chlorobiphenyls and bromobisphenols were extracted by liquid-liquid extraction and were quantified by using the GC equipped with an ECD as reported previously ^[48, 61]^. Chloroethenes were analyzed using GC with a flame ionization detector and a Gas-Pro column (30 m × 0.32 mm; Agilent J&W Scientific, Folsom, CA, USA) ^[33]^. MeHg was detected using high-performance liquid chromatography (HPLC) coupled with inductively coupled plasma mass spectrometer (ICP-MS, iCAP Q, Thermo Fisher Scietific). Chromatographic separation was achieved on a PLATISIL ODS column (150 × 4.6 mm, 5 µm). The ICP-MS was operated under the following conditions: RF power 1550 W; plasma gas flow rate 14 L/min; auxiliary gas flow rate 0.8 L/min; nebulizer gas flow rate 0.85 L/min; collision gas flow rate 0 mL/min (standard mode); sampling depth 5 mm; and dwell time 200 ms. The detection limit was around 0.04 µg/L MeHg. Acetate was detected using GC equipped with FlD (Agilent, Wilmington, DE, USA) and a DB-FFAP column (30 m × 0.25 mm × 0.25 μm; Agilent) ^[62]^. Formate was detected using ion chromatography (ICS-600, Thermo Scientific, USA). Carbon dioxide was determined according to protocol specified in ASTM D513-16 (2024)^[63]^, with uninoculated medium serving as a reference control.

### Global-scale marine and terrestrial metagenomic datasets collection and assembly

Metagenomes of marine and terrestrial settings were downloaded from the National Center for Biotechnology Information (NCBI) Sequence Read Archive (SRA) database (NCBI-SRA) (Supplementary Data 3; Supplementary Data 4). Metagenomic raw reads data were filtered to remove low-quality bases/reads using trim_galore^[64]^ (v0.6.10) with default parameters. Clean reads from each sample were assembled individually into contigs using MEGAHIT v1.2.9 (k-mer: 21,29,39,59,79,99,119,141)^[65]^ with subsequent quality assessment using QUAST^[66]^ (v5.0.2). Contigs were taxonomically assigned to taxa using both CAT^[67]^ (v5.2.3) and Kaiju^[68]^ (v1.9.2) to improve the accuracy and breadth of annotations. The contigs were annotated with CAT by predicting open reading frames (ORFs) with Prodigal^[69]^ (v2.6.3; parameter: -meta) and by comparing them with DIAMOND blastp^[70]^ (v2.1.7) to the non-redundant set of proteins in GTDB (GTDB taxonomy release_214)^[71]^. In addition, the contigs annotation with Kaiju was performed utilizing the NCBI nr database that included bacteria, archaea, viruses, fungi, and other eukaryotic microorganisms for annotating contigs with default parameters.

### Identification of *hgcAB* and abundance calculation

To identify the *hgcAB* gene cluster, ORFs of contigs were predicted using Prodigal^[69]^ (v2.6.3; parameter: -meta), with which *hgcAB* were identified based on both protein sequence similarity and conserved regions/motifs against Hg-MATE-Db^[72]^. In brief, protein sequences were searched against Hg-MATE-Db using hmmsearch with score ≥300 or e-value ≤ 1e-50 as the cutoff. Results were further confirmed by examining the presence of conserved motifs (N(V/I)WCA(A/G)) in *hgcA* and (CX2CX2CX3C) in *hgcB*, respectively^[8, 10]^. To determine the relative abundance of *hgcAB* gene clusters in contigs, clean reads were mapped to the contigs using CoverM (v 0.6.0, https://github.com/wwood/CoverM/) with parameter ‘-contig’ and cut-off values of 75% identity and 75% alignment coverage for mapped reads, which generated coverage profiles of each contig and normalized as RPKM. The RPKM were calculated using the equation:

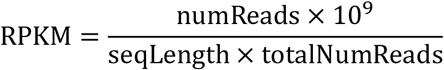

Where numReads is the number of reads mapped to a sequence; seqLength is the length of the sequence; totalNumReads is the total number of mapped reads of a sample. Then, the relative abundance of *hgcAB* gene clusters was calculated by dividing RPKM values of individual genes by the sum of RPKM values of all genes.

### Correlation analysis between *hgcAB* abundance and halomethane concentrations

Halomethanes data was collected from the HalOcAt (Halocarbons in the Ocean and Atmosphere) database (https://halocat.geomar.de/)^[73]^and Atom (Atmospheric Tomography) database (https://doi.org/10.3334/ORNLDAAC/1925) ^[74]^. All available halomethane species (including chloromethanes, bromomethanes, and mixed halomethanes) from both datasets were retrieved. A spatial grid-based correlation analysis was conducted to investigate the relationship between *hgcAB* gene abundance and halomethane concentrations. The global map was divided into grid cells with a resolution of 5° in both longitude and latitude. For each grid cell (5°×5°), the average abundance of *hgcAB* genes and the average concentration of halomethanes were calculated from all data points falling within that cell. Only grid cells containing both types of data were retained for analysis; cells with only one type of data or none were excluded. A correlation analysis was then performed on the paired average values from the selected grid cells to assess the relationship between the two variables.

### Statistical analysis

Correlation analyses were performed using Pearson’s correlation coefficient in IBM SPSS Statistics 26 software. Significance levels were set at *p <* 0.05, and all statistical tests were two-tailed.

## Acknowledgements

This study was financially supported by the National Key Research and Development Program of China (2024YFC3713303, 2024YFC3713300) and the National Natural Science Foundation of China (42177001).

## References

1. Dastoor A, Angot H, Bieser J, et al. Arctic mercury cycling. Nat Rev Earth Env, 2022. 3(4): p. 270–286. 10.1038/s43017-022-00269-w

2. Jonsson S, Skyllberg U, Nilsson M B, et al. Differentiated availability of geochemical mercury pools controls methylmercury levels in estuarine sediment and biota. Nat Commun, 2014. 5(1): p. 4624. 10.1038/ncomms5624

3. Lei P, Zhang J, Yu R Q, et al. Elevated atmospheric CO2 decreases methylmercury production in freshwater lakes. Nat Commun, 2025. 17(1): p. 1037. 10.1038/s41467-025-67788-0

4. Motta L C, Blum J D, Popp B N, et al. Mercury isotopic evidence for the importance of particles as a source of mercury to marine organisms. Proc Natl Acad Sci U S A, 2022. 119(44): p. e2208183119. 10.1073/pnas.2208183119

5. Stern A H, Smith A E. An assessment of the cord blood:maternal blood methylmercury ratio: implications for risk assessment. Environ Health Perspect, 2003. 111(12): p. 1465–70. 10.1289/ehp.6187

6. Axelrad D A, Bellinger D C, Ryan L M, et al. Dose-response relationship of prenatal mercury exposure and IQ: an integrative analysis of epidemiologic data. Environ Health Perspect, 2007. 115(4): p. 609–15. 10.1289/ehp.9303

7. Roman H A, Walsh T L, Coull B A, et al. Evaluation of the cardiovascular effects of methylmercury exposures: current evidence supports development of a dose-response function for regulatory benefits analysis. Environ Health Perspect, 2011. 119(5): p. 607–14. 10.1289/ehp.1003012

8. Parks J M, Johs A, Podar M, et al. The genetic basis for bacterial mercury methylation. Science, 2013. 339(6125): p. 1332–5. 10.1126/science.1230667

9. Podar M, Gilmour C C, Brandt C C, et al. Global prevalence and distribution of genes and microorganisms involved in mercury methylation. Sci Adv, 2015. 1(9): p. e1500675. 10.1126/sciadv.1500675

10. Li Z, Wei T, He L, et al. Genomic potential for mercury biotransformation in marine sediments across marginal slope to hadal zone. Nat Commun, 2025. 16(1): p. 8655. 10.1038/s41467-025-63808-1

11. Gilmour C C, Podar M, Bullock A L, et al. Mercury methylation by novel microorganisms from new environments. Environ Sci Technol, 2013. 47(20): p. 11810–20. 10.1021/es403075t

12. Qian C, Chen H, Johs A, et al. Quantitative Proteomic Analysis of Biological Processes and Responses of the Bacterium *Desulfovibrio desulfuricans* ND132 upon Deletion of Its Mercury Methylation Genes. Proteomics, 2018. 18(17): p. e1700479. 10.1002/pmic.201700479

13. Zhang Z, Zhang C, Chen W, et al. Antibiotic Pollution Elevates Microbial Methylmercury Production. Environ Sci Technol, 2026. 60(13): p. 10274–10284. 10.1021/acs.est.6c01533

14. McDaniel Elizabeth A, Peterson Benjamin D, Stevens Sarah L R, et al. Expanded Phylogenetic Diversity and Metabolic Flexibility of Mercury-Methylating Microorganisms. mSystems, 2020. 5(4): p. 10.1128/msystems.00299-20. http://doi.org/10.1128/msystems.00299-20

15. Gionfriddo Caitlin M, Soren Ally B, Wymore Ann M, et al. Transcriptional Control of hgcAB by an ArsR-Like Regulator in Pseudodesulfovibrio mercurii ND132. Applied and Environmental Microbiology, 2023. 89(4): p. e01768–22. 10.1128/aem.01768-22

16. Hillesland K L, Lim S, Flowers J J, et al. Erosion of functional independence early in the evolution of a microbial mutualism. Proc Natl Acad Sci U S A, 2014. 111(41): p. 14822–7. 10.1073/pnas.1407986111

17. Wu B, Liu F, Fang W, et al. Microbial sulfur metabolism and environmental implications. Sci Total Environ, 2021. 778: p. 146085. 10.1016/j.scitotenv.2021.146085

18. Cooper C J, Zheng K, Rush K W, et al. Structure determination of the HgcAB complex using metagenome sequence data: insights into microbial mercury methylation. Commun Biol, 2020. 3(1): p. 320. 10.1038/s42003-020-1047-5

19. Bridwell-Rabb J, Drennan C L. Vitamin B(12) in the spotlight again. Curr Opin Chem Biol, 2017. 37: p. 63–70. 10.1016/j.cbpa.2017.01.013

20. Soder-Walz J M, Deobald D, Vicent T, et al. MecE, MecB, and MecC proteins orchestrate methyl group transfer during dichloromethane fermentation. Appl Environ Microbiol, 2024. 90(10): p. e0097824. 10.1128/aem.00978-24

21. Murdoch R W, Chen G, Kara Murdoch F, et al. Identification and widespread environmental distribution of a gene cassette implicated in anaerobic dichloromethane degradation. Glob Chang Biol, 2022. 28(7): p. 2396–2412. 10.1111/gcb.16068

22. Obrist D, Tas E, Peleg M, et al. Bromine-induced oxidation of mercury in the mid-latitude atmosphere. Nat Geosci, 2011. 4(1): p. 22–26. 10.1038/ngeo1018

23. Zerkle A L, Claire M W, Rocco T D, et al. Sulfur and mercury MIF suggest volcanic contributions to Earth’s atmosphere at 2.7 Ga. Geochem Persp Let, 2021. 18: p. 48–52. 10.7185/geochemlet.2124

24. McConnell J C, Henderson G S, Barrie L, et al. Photochemical bromine production implicated in Arctic boundary-layer ozone depletion. Nature, 1992. 355(6356): p. 150–152. 10.1038/355150a0

25. Gilmour C C, Elias D A, Kucken A M, et al. Sulfate-reducing bacterium *Desulfovibrio desulfuricans* ND132 as a model for understanding bacterial mercury methylation. Appl Environ Microbiol, 2011. 77(12): p. 3938–51. 10.1128/AEM.02993-10

26. Hu H, Lin H, Zheng W, et al. Oxidation and methylation of dissolved elemental mercury by anaerobic bacteria. Nat Geosci, 2013. 6(9): p. 751–754. 10.1038/ngeo1894

27. Brown Steven D, Gilmour Cynthia C, Kucken Amy M, et al. Genome Sequence of the Mercury-Methylating Strain Desulfovibrio desulfuricans ND132. J Bacteriol, 2011. 193(8): p. 2078–2079. 10.1128/jb.00170-11

28. Methé B A, Nelson K E, Eisen J A, et al. Genome of *Geobacter sulfurreducens*: Metal Reduction in Subsurface Environments. Science, 2003. 302(5652): p. 1967–1969. 10.1126/science.1088727

29. Wang S, Chng K R, Wilm A, et al. Genomic characterization of three unique *Dehalococcoides* that respire on persistent polychlorinated biphenyls. Proc Natl Acad Sci U S A, 2014. 111(33): p. 12103–8. 10.1073/pnas.1404845111

30. Chen G, Fisch A R, Gibson C M, et al. Mineralization versus fermentation: evidence for two distinct anaerobic bacterial degradation pathways for dichloromethane. ISME J, 2020. 14(4): p. 959–970. 10.1038/s41396-019-0579-5

31. Mägli A, Wendt M, Leisinger T. Isolation and characterization of *Dehalobacterium formicoaceticum* gen. nov. sp. nov., a strictly anaerobic bacterium utilizing dichloromethane as source of carbon and energy. Arch Microbiol, 1996. 166(2): p. 101–108. 10.1007/s002030050362

32. Chen G, Murdoch Robert W, Mack E E, et al. Complete Genome Sequence of *Dehalobacterium formicoaceticum* Strain DMC, a Strictly Anaerobic Dichloromethane-Degrading Bacterium. Genome Announc, 2017. 5(37): p. 10.1128/genomea.00897-17. http://doi.org/10.1128/genomea.00897-17

33. Zhou N, Li Q, Liang Z, et al. Microbially-mediated halogenation and dehalogenation cycling of organohalides in the ocean. Nat Commun, 2025. 16(1): p. 10670. 10.1038/s41467-025-65696-x

34. Zhang Y, Zhang P, Song Z, et al. An updated global mercury budget from a coupled atmosphere-land-ocean model: 40% more re-emissions buffer the effect of primary emission reductions. One Earth, 2023. 6(3): p. 316–325. 10.1016/j.oneear.2023.02.004

35. World Meteorological Organization. Update on Ozone-Depleting Substances (ODSs) and Other Gases of Interest to the Montreal Protocol.Scientific Assessment of Ozone Depletion: 2022 World Meteorological Organization (WMO): Geneva. p. 1–112. 2022.

36. Wu Z, Li Z, Shao B, et al. Impact of dissolved organic matter and environmental factors on methylmercury concentrations across aquatic ecosystems inferred from a global dataset. Chemosphere, 2022. 294: p. 133713. 10.1016/j.chemosphere.2022.133713

37. Semeniuk K, Dastoor A. Development of a global ocean mercury model with a methylation cycle: Outstanding issues. Global Biogeochem Cycles, 2017. 31(2): p. 400–433. 10.1002/2016GB005452

38. Zhang Y, Soerensen A L, Schartup A T, et al. A Global Model for Methylmercury Formation and Uptake at the Base of Marine Food Webs. Global Biogeochem Cycles, 2020. 34(2): p. e2019GB006348. 10.1029/2019GB006348

39. Bowman K L, Lamborg C H, Agather A M. A global perspective on mercury cycling in the ocean. Sci Total Environ, 2020. 710: p. 136166. 10.1016/j.scitotenv.2019.136166

40. Grigal D F. Mercury Sequestration in Forests and Peatlands. J Environ Qual, 2003. 32(2): p. 393–405. 10.2134/jeq2003.3930

41. Selvendiran P, Driscoll C T, Bushey J T, et al. Wetland influence on mercury fate and transport in a temperate forested watershed. Environ Pollut, 2008. 154(1): p. 46–55. 10.1016/j.envpol.2007.12.005

42. Fleck J A, Marvin-DiPasquale M, Eagles-Smith C A, et al. Mercury and methylmercury in aquatic sediment across western North America. Sci Total Environ, 2016. 568: p. 727–738. 10.1016/j.scitotenv.2016.03.044

43. Lin H, Moody E R R, Williams T A, et al. On the Origin and Evolution of Microbial Mercury Methylation. Genome Biol Evol, 2023. 15(4). 10.1093/gbe/evad051

44. Fincker M, Spormann A M. Biochemistry of catabolic reductive dehalogenation. Annu Rev Biochem, 2017. 86: p. 357–386. 10.1146/annurev-biochem-061516-044829

45. Bommer M, Kunze C, Fesseler J, et al. Structural basis for organohalide respiration. Science, 2014. 346(6208): p. 455–8. 10.1126/science.1258118

46. Payne K A P, Quezada C P, Fisher K, et al. Reductive dehalogenase structure suggests a mechanism for B12-dependent dehalogenation. Nature, 2015. 517(7535): p. 513–516. 10.1038/nature13901

47. Xu G, Zhao S, Rogers M J, et al. Global prevalence of organohalide-respiring bacteria dechlorinating polychlorinated biphenyls in sewage sludge. Microbiome, 2024. 12(1): p. 54. 10.1186/s40168-024-01754-8

48. Qiu L, Fang W, He H, et al. Organohalide-Respiring Bacteria in Polluted Urban Rivers Employ Novel Bifunctional Reductive Dehalogenases to Dechlorinate Polychlorinated Biphenyls and Tetrachloroethene. Environ Sci Technol, 2020. 54(14): p. 8791–8800. 10.1021/acs.est.0c01569

49. Han Y, Deng Z, Peng Y, et al. Evidence of microbial reductive dehalogenation in deep-sea cold seeps and its implications for biogeochemical cycles. Microbiome, 2025. 13(1): p. 156. 10.1186/s40168-025-02147-1

50. Gilmour C C, Tuttle J H, Means J C. Anaerobic microbial methylation of inorganic tin in estuarine sediment slurries. Microb Ecol, 1987. 14(3): p. 233–42. 10.1007/bf02012943

51. Caccavo F, Jr., Lonergan D J, Lovley D R, et al. *Geobacter sulfurreducens* sp. nov., a hydrogen- and acetate-oxidizing dissimilatory metal-reducing microorganism. Appl Environ Microbiol, 1994. 60(10): p. 3752–9. 10.1128/aem.60.10.3752-3759.1994

52. Zhang Z, Si R, Lv J, et al. Effects of Extracellular Polymeric Substances on the Formation and Methylation of Mercury Sulfide Nanoparticles. Environ Sci Technol, 2020. 54(13): p. 8061–8071. 10.1021/acs.est.0c01456

53. Lu Y, Zhang S, Liu Q, et al. Nitrobenzene reduction promoted by the integration of carbon nanotubes and Geobacter sulfurreducens. Environ Pollut, 2023. 325: p. 121444. 10.1016/j.envpol.2023.121444

54. Jia B, Shen R, Li Z, et al. Synergistic integration of in situ generated Fe-S species and organohalide-respiring bacteria for complete dechlorination of carbon tetrachloride in groundwater. unpublished.

55. Gilmour C C, Soren A B, Gionfriddo C M, et al. *Pseudodesulfovibrio mercurii* sp. nov., a mercury-methylating bacterium isolated from sediment. Int J Syst Evol Microbiol, 2019. 71(3). 10.1099/ijsem.0.004697

56. Mägli A, Messmer M, Leisinger T. Metabolism of Dichloromethane by the Strict Anaerobe *Dehalobacterium formicoaceticum*. Appl Environ Microbiol, 1998. 64(2): p. 646–650. 10.1128/AEM.64.2.646-650.1998

57. Zheng K, Rush K W, Date S S, et al. S-adenosyl-L-methionine is the unexpected methyl donor for the methylation of mercury by the membrane-associated HgcAB complex. Proc Natl Acad Sci U S A, 2024. 121(47): p. e2408086121. 10.1073/pnas.2408086121

58. Shen R, Liang Z, Lu Q, et al. Spatiotemporal profiling and succession of microbial communities in landfills based on a cross-kingdom abundance quantification method. Water Res, 2025. 277: p. 123334. 10.1016/j.watres.2025.123334

59. Wang Y, Qian P Y. Conservative fragments in bacterial 16S rRNA genes and primer design for 16S ribosomal DNA amplicons in metagenomic studies. PLoS One, 2009. 4(10): p. e7401. 10.1371/journal.pone.0007401

60. Callahan B J, McMurdie P J, Rosen M J, et al. DADA2: High-resolution sample inference from Illumina amplicon data. Nat Methods, 2016. 13(7): p. 581–3. 10.1038/nmeth.3869

61. Xu G, Ng H L, Chen C, et al. Combatting multiple aromatic organohalide pollutants in sediments by bioaugmentation with a single Dehalococcoides. Water Res, 2024. 255: p. 121447. 10.1016/j.watres.2024.121447

62. Weber T E, Trabue S L, Ziemer C J, et al. Evaluation of elevated dietary corn fiber from corn germ meal in growing female pigs. J Anim Sci, 2010. 88(1): p. 192–201. 10.2527/jas.2009-1896

63. ASTM International, Standard Test Methods for Total and Dissolved Carbon Dioxide in Water. 2024.

64. Martin M. Cutadapt removes adapter sequences from high-throughput sequencing reads. EMBnet. j, 2011. 17: p. 10–12.

65. Li D, Liu C M, Luo R, et al. MEGAHIT: an ultra-fast single-node solution for large and complex metagenomics assembly via succinct de Bruijn graph. Bioinformatics, 2015. 31(10): p. 1674–6. 10.1093/bioinformatics/btv033

66. Quast C, Pruesse E, Yilmaz P, et al. The SILVA ribosomal RNA gene database project: improved data processing and web-based tools. Nucleic Acids Res, 2013. 41(Database issue): p. D590–6. 10.1093/nar/gks1219

67. von Meijenfeldt F A B, Arkhipova K, Cambuy D D, et al. Robust taxonomic classification of uncharted microbial sequences and bins with CAT and BAT. Genome Biol, 2019. 20(1): p. 217. 10.1186/s13059-019-1817-x

68. Menzel P, Ng K L, Krogh A. Fast and sensitive taxonomic classification for metagenomics with Kaiju. Nat Commun, 2016. 7: p. 11257. 10.1038/ncomms11257

69. Hyatt D, Chen G L, Locascio P F, et al. Prodigal: prokaryotic gene recognition and translation initiation site identification. BMC Bioinform, 2010. 11: p. 119. 10.1186/1471-2105-11-119

70. Buchfink B, Xie C, Huson D H. Fast and sensitive protein alignment using DIAMOND. Nat Methods, 2015. 12(1): p. 59–60. 10.1038/nmeth.3176

71. Parks D H, Chuvochina M, Rinke C, et al. GTDB: an ongoing census of bacterial and archaeal diversity through a phylogenetically consistent, rank normalized and complete genome-based taxonomy. Nucleic Acids Res, 2022. 50(D1): p. D785–d794. 10.1093/nar/gkab776

72. Capo E, Peterson B D, Kim M, et al. A consensus protocol for the recovery of mercury methylation genes from metagenomes. Mol Ecol Resour, 2023. 23(1): p. 190–204. 10.1111/1755-0998.13687

73. Quack B, Archer S, Artuso F, et al., The HalOcAt-database of volatile short-lived halogenated hydrocarbons in ocean and atmosphere (1990-2018). 2025, PANGAEA.

74. Wofsy S C, Afshar S, Allen H M, et al., ATom: Merged Atmospheric Chemistry, Trace Gases, and Aerosols, Version 2. 2021.

