## Supplementary material for "Metabolic dehalogenation of halomethanes by mercury methylators": Fig. S1-S10

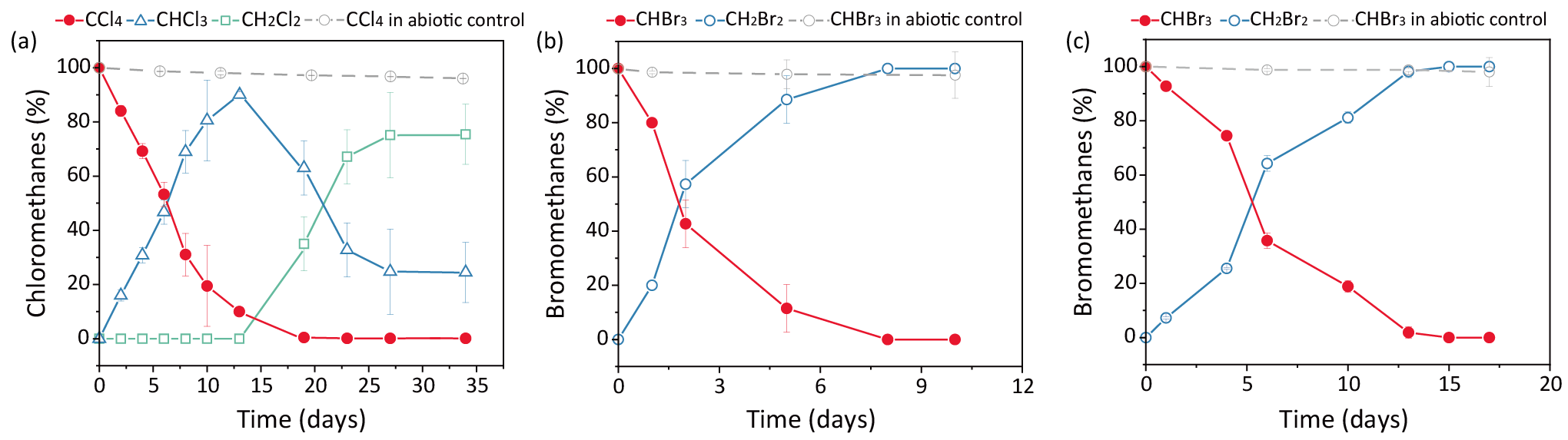

**Figure S1. Reductive dehalogenation in pre-grown pure cultures of *P. mercurii* ND132 and *G. sulfurreducens* PCA.** (a) Kinetics of CCl_4_ dechlorination in *G. sulfurreducens* PCA and its abiotic control; Kinetics of bromoform debromination in pre-grown pure cultures of (b) *P. mercurii* ND132 and (c) *G. sulfurreducens* PCA and their abiotic controls. Error bars represent SDs of triplicate cultures.

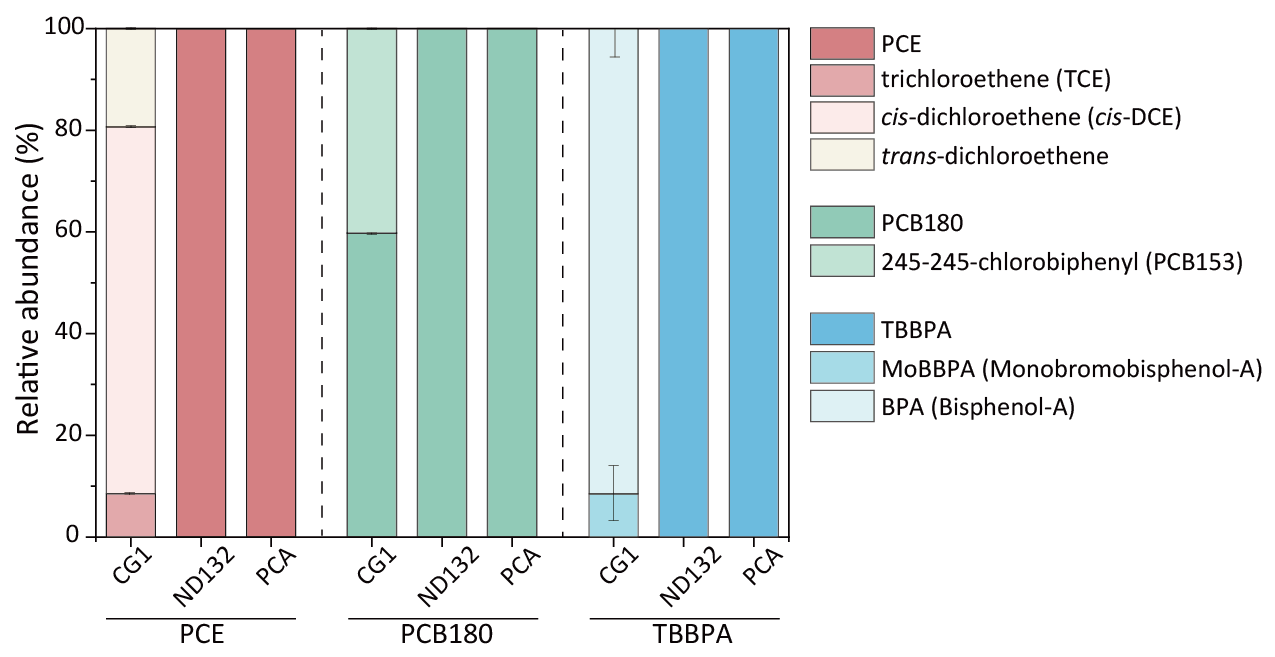

**Figure S2.** Dehalogenation of perchloroethene (PCE), 2345-245-chlorobiphenyl (PCB180) or tetrabromobisphenol-A (TBBPA) in pre-grown pure cultures of *P. mercurii* ND132 and *G. sulfurreducens* PCA after one month of incubation. *D. mccartyi* CG1 was employed as a positive control. Error bars represent SDs of triplicate cultures.

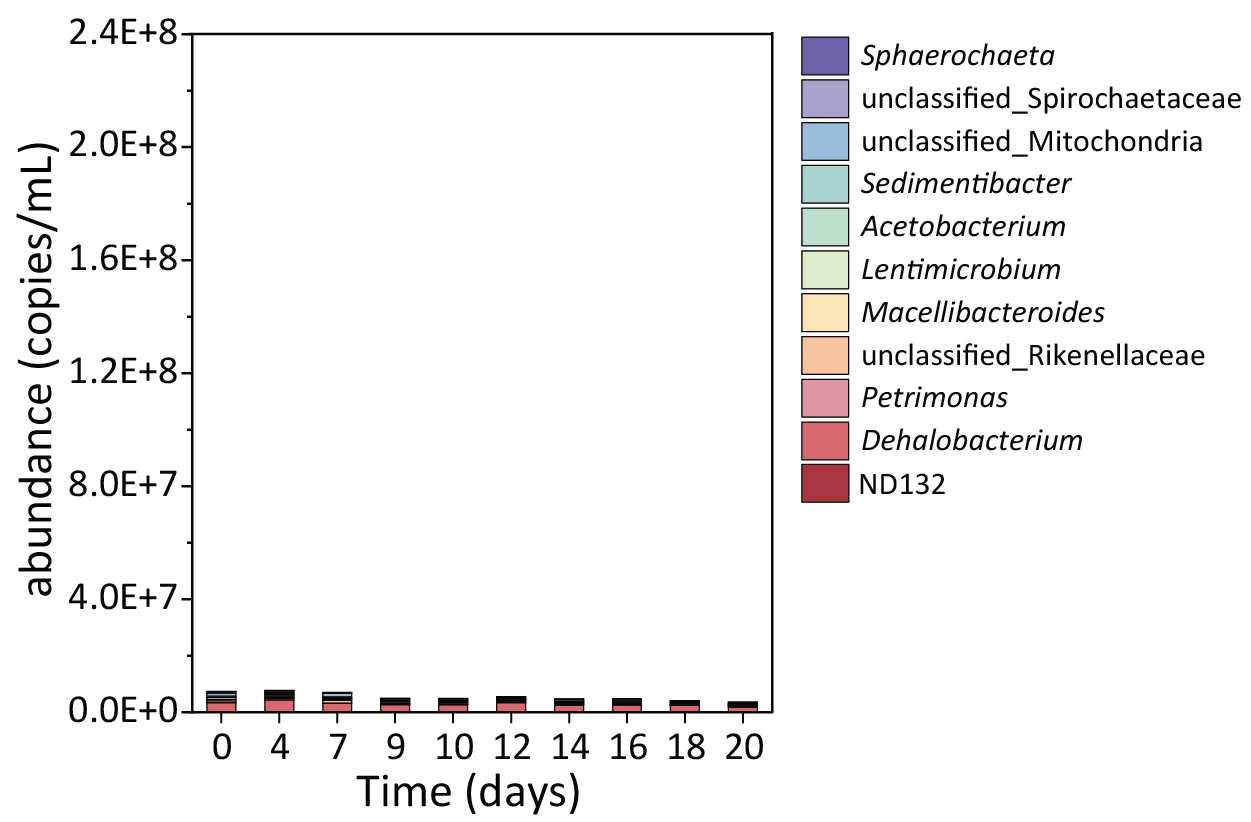

**Figure S3.** Temporal changes in microbial community composition at the genus level in biotic control cultures without amendment of chloromethanes.

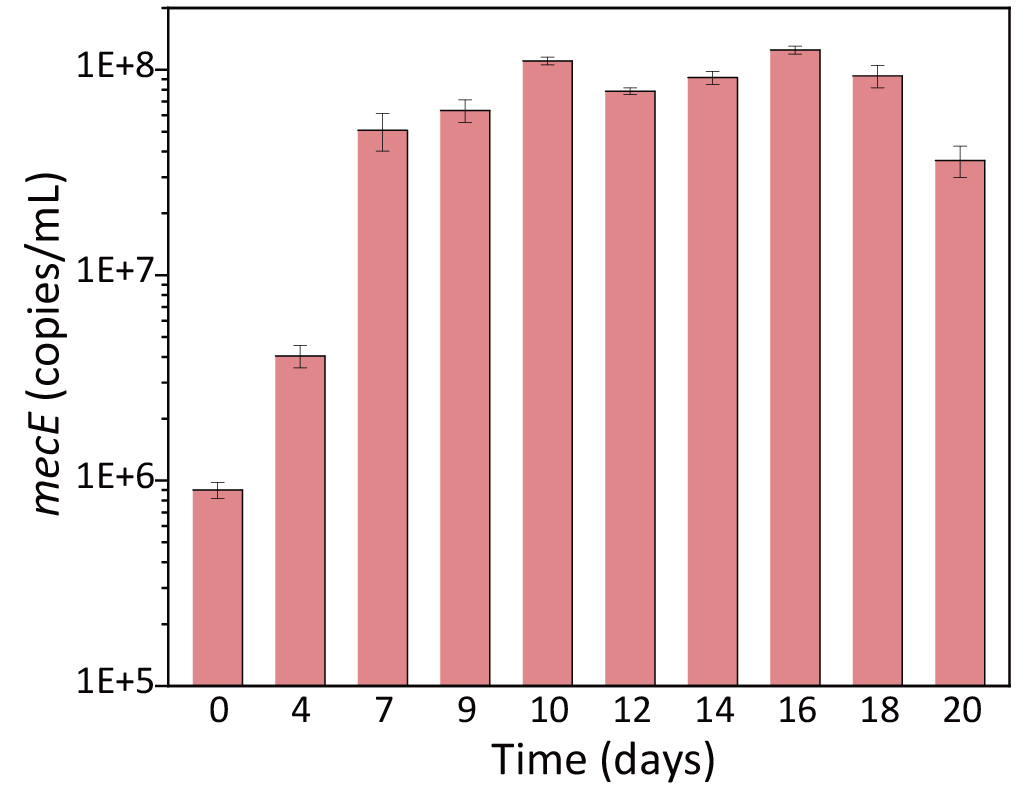

**Figure S4.** The qPCR quantification of the *mecE* gene in the synthetic microbial consortium.

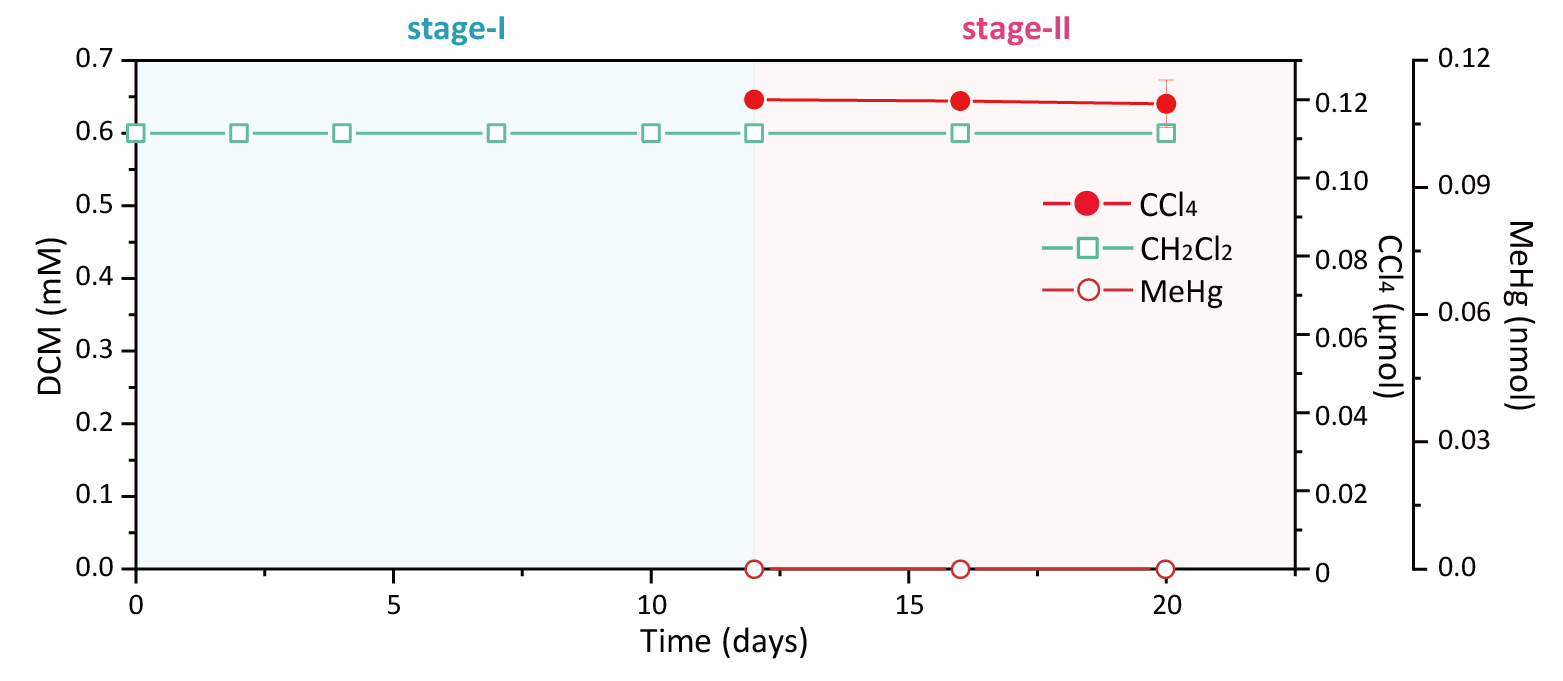

**Figure S5.** Kinetics of chloromethanes dechlorination/degradation and Hg methylation in the abiotic control culture.

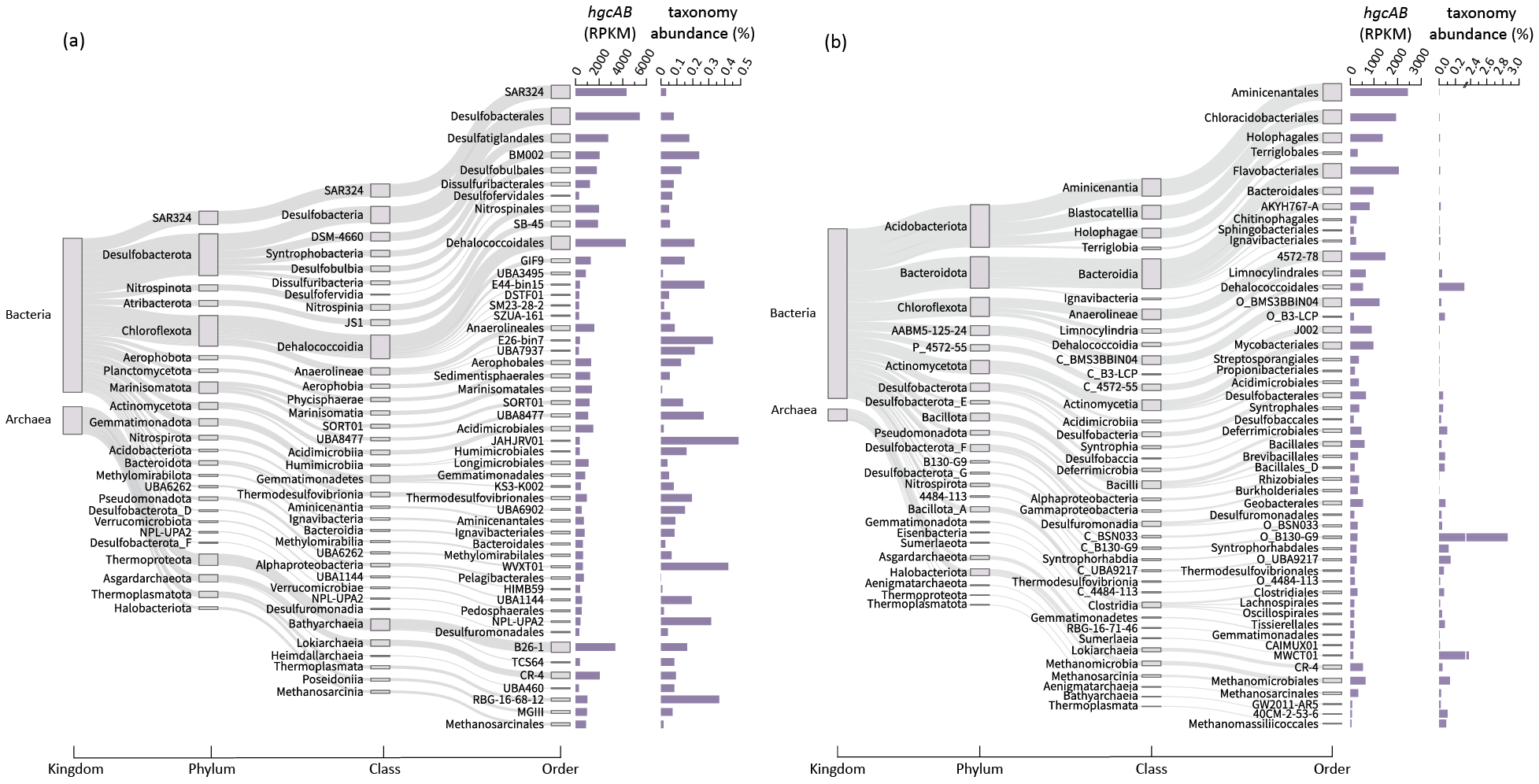

**Figure S6**. Major taxonomic distribution of *hgcAB* genes in prokaryotes in marine (a) and terrestrial (b) metagenomes.

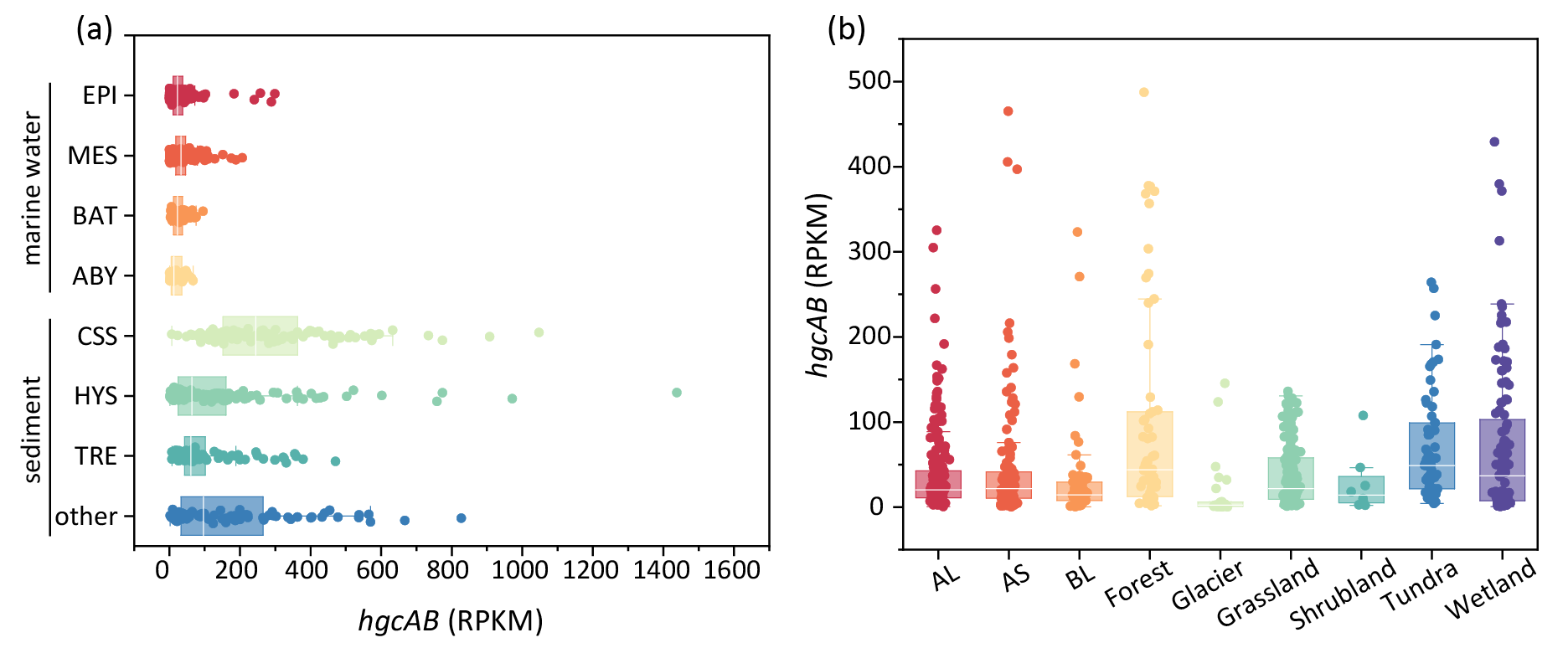

**Figure S7**. Global geographic distribution of *hgcAB* genes in prokaryotes. (a) Marine environments. (b) Terrestrial environments. EPI, epipelagic layer; MES, mesopelagic layer; BAT, bathypelagic layer; ABY, abyssopelagic layer; CSS, cold seep sediment; HVS, hydrothermal vent sediment; TRE, trench sediment; AL, Agricultural Land; AS, Artificial Surfaces; BL, Bare Land.

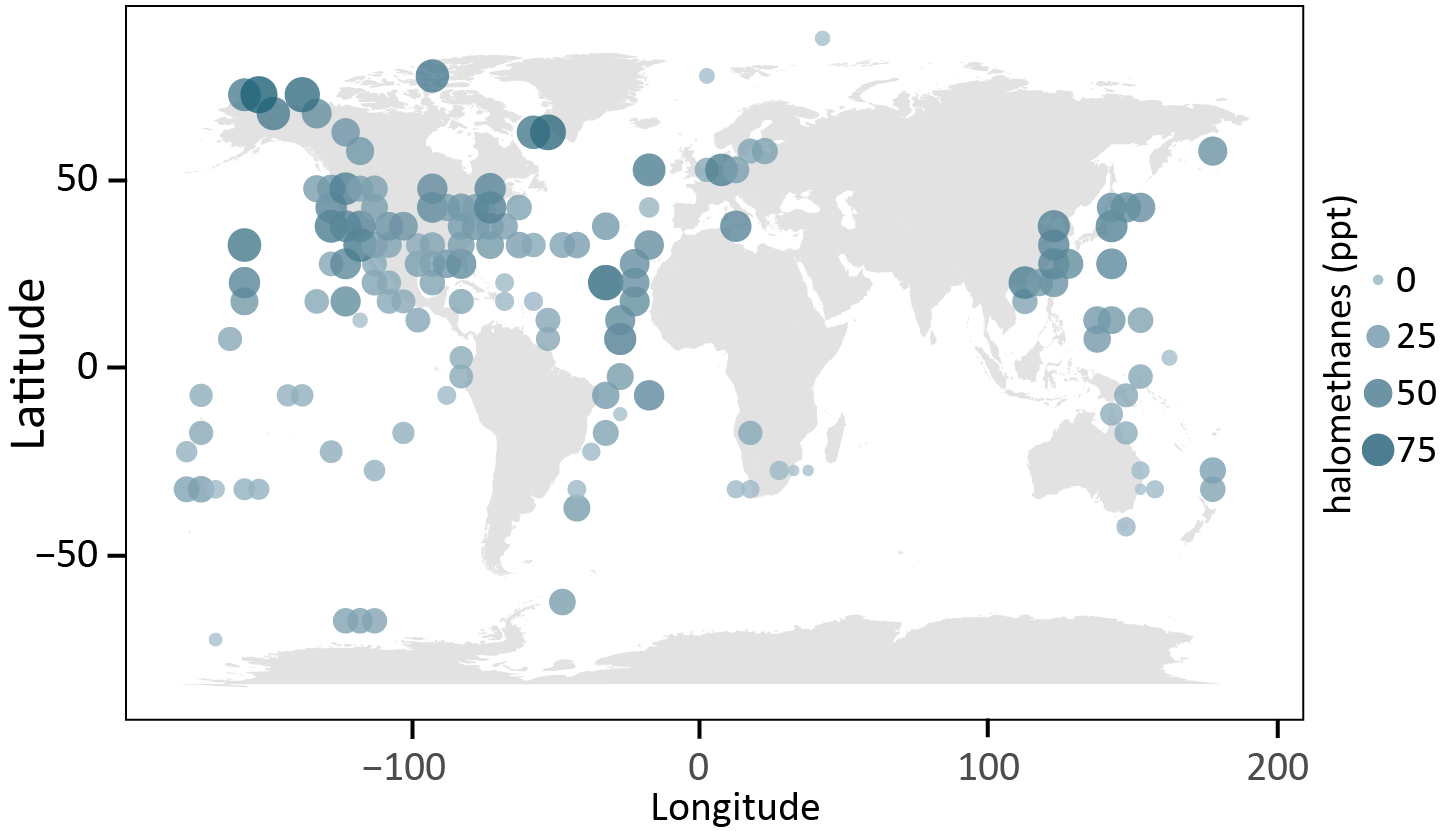

**Figure S8.** Global geographic distribution of atmospheric halomethanes. Values represent the average concentration of halomethanes within each 5°×5° (longitude × latitude) grid cell. Only grid cells containing detectable *hgcAB* genes are shown.

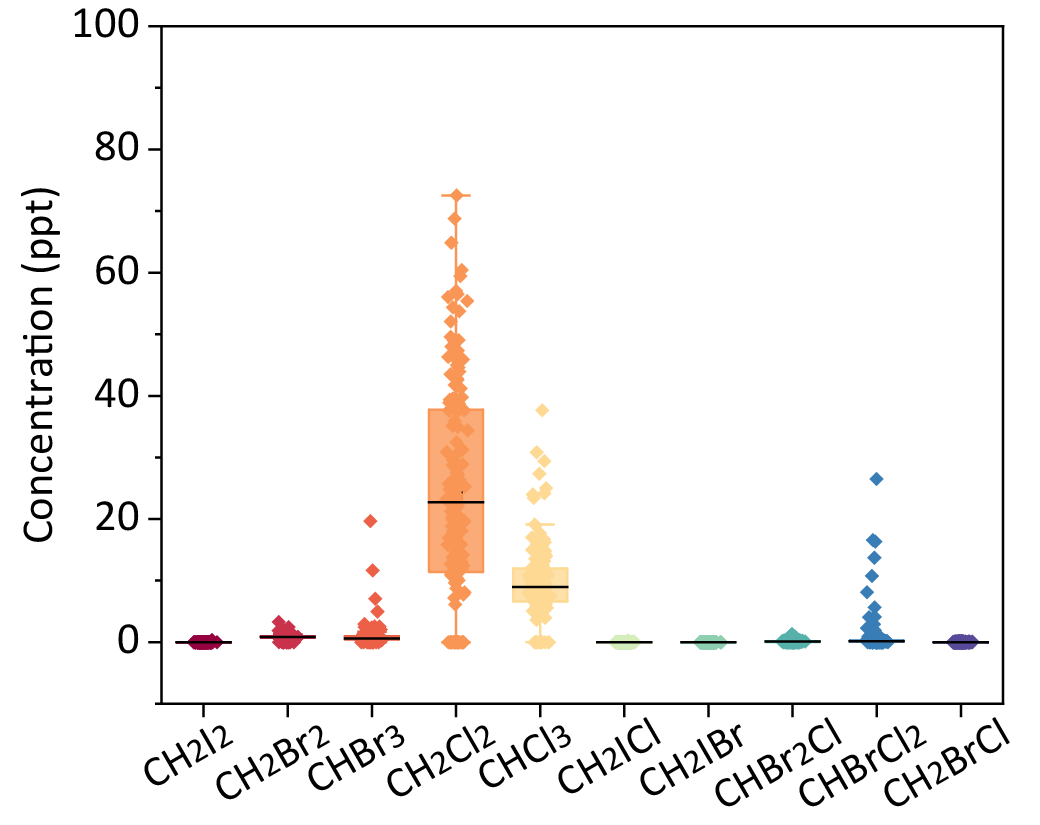

**Figure S9.** Halomethanes and their respective concentrations included in the correlation analysis.

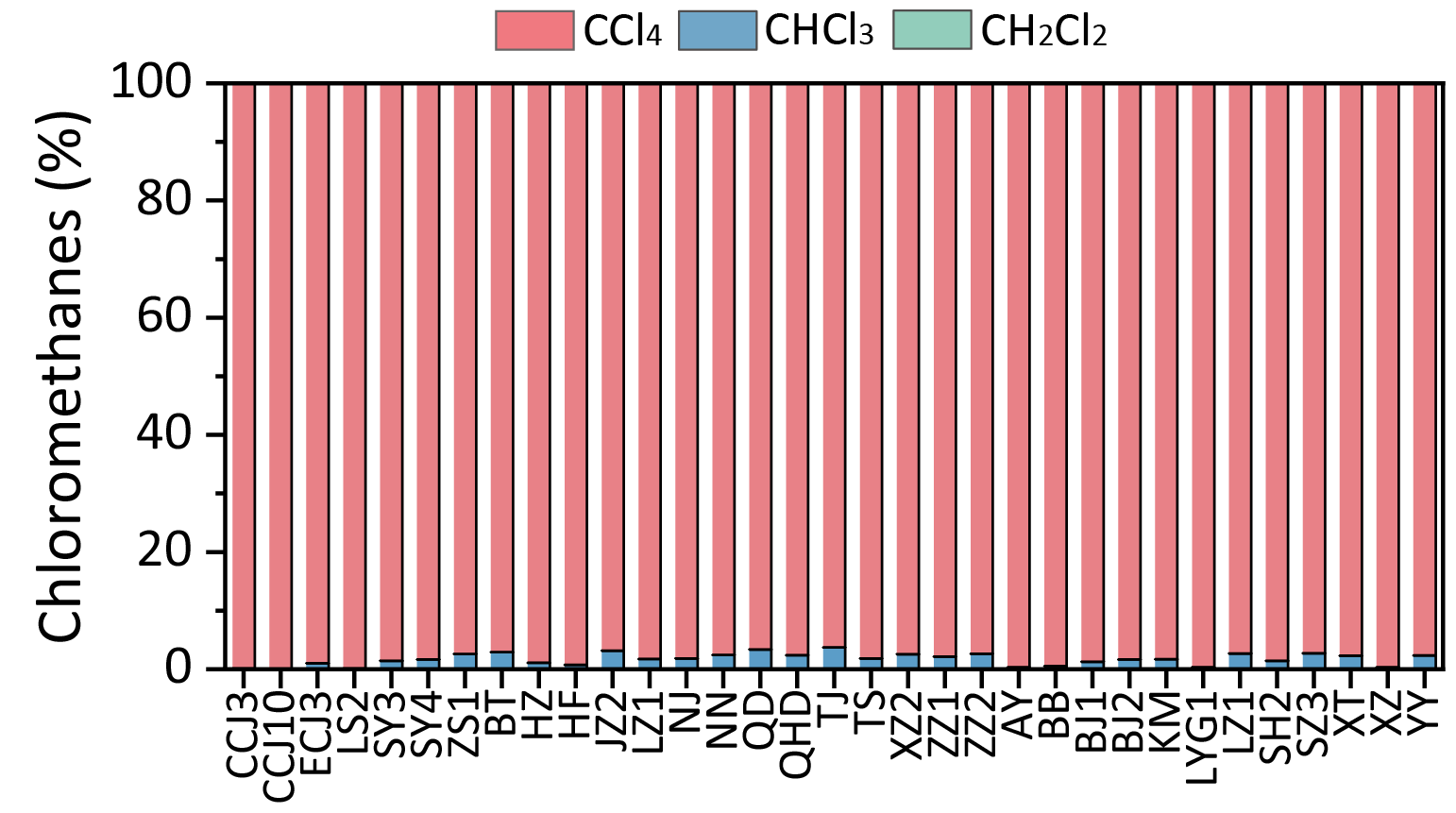

**Figure S10.** Reductive dechlorination of CCl_4_ in autoclaved microcosms (abiotic controls) after 7 days incubation.
